# Nutritional Programming Improves Dietary Plant Protein Utilization in Zebrafish *Danio rerio*

**DOI:** 10.1101/846725

**Authors:** Karolina Kwasek, Michal Wojno, Genciana Terova, Vance J. McCracken, Giovanni S. Molinari, Federica Iannini

**Author notes:** Corresponding author: Center for Fisheries, Aquaculture, and Aquatic Sciences, Southern Illinois University, 1125 Lincoln Drive, Life Science II room 175, Carbondale, IL 62901.

## Abstract

Nutritional Programming (NP) has been shown to counteract the negative effects of dietary plant protein (PP) by introducing PP at an early age towards enhancement of PP utilization during later life stages. This study explored the effect of NP and its induction time on growth, expression of appetite-stimulating hormones, and any morphological changes in the gut possibly responsible for improved dietary PP utilization. At 3 days post-hatch (dph) zebrafish were distributed into 12 (3 L) tanks, 100 larvae per tank. This study included four groups: 1) The control (NP-FM) group received fishmeal (FM)-based diet from 13-36 dph and was challenged with PP-based diet during 36-66 dph; 2) The NP-PP group received NP with dietary PP in larval stage via live food enrichment during 3-13 dph followed by FM diet during 13-36 dph and PP diet during 36-66 dph; 3) The T-NP group received NP between 13-23 dph through PP diet followed by FM diet during 23-36 dph and PP diet during 36-66 dph; and 4) The PP group received PP diet from 13-66 dph. During the PP challenge the T-NP group achieved the highest weight gain compared to control and PP. *Ghrelin* expression in the brain was higher in T-NP compared to NP-FM and NP-PP, while in the gut it was reduced in both NP-PP and T-NP groups. *Cholecystokinin* expression showed an opposite trend to *ghrelin*. The brain *neuropeptide Y* expression was lower in NP-PP compared to PP but not different with NP-FM and T-NP groups. The highest villus length to width ratio in the middle intestine was found in T-NP compared to all other groups. The study suggests that NP induced during juveniles stages improves zebrafish growth and affects digestive hormone regulation and morphology of the intestinal lining – possible mechanisms behind the improved PP utilization in pre-adult zebrafish stages.

## Introduction

Replacement of fishmeal (FM) in fish diets with plant protein (PP) has been an ongoing challenge in the aquaculture industry. High-quality PP sources such as soy or pea protein concentrates and wheat or corn gluten have been widely used by the feed industry since their digestibility in some species is comparable to FM. However, their price can often exceed the cost of marine raw materials. Although some progress with utilization of lower-quality PP, such as soybean meal, has been made, a number of concerns must still be overcome including low palatability, imbalanced amino acid profile, and a presence of anti-nutritional factors responsible for inducing intestinal inflammation, to maintain acceptable growth rates and feed efficiency values at high FM substitution levels. Thus, the aquafeed industry has focused on ways to include some of the more cost-effective alternative sources of protein that will not only help to further replace FM but also substitute some of the expensive high-quality PP concentrates and provide more flexibility in feed formulations using a wider range of available raw materials.

Nutritional programming (NP) has been found to be a promising approach to counteract the negative effects of PP in feeds. NP is a way of modifying or inducing specific physiological responses during vulnerable early developmental stages of an animal that will help to cope with specific (environmental) challenges in the animal later in (adult) life. NP in fish only recently has received more attention and some studies already indicate it is possible to “program” fish during their young age with alternative raw materials or nutrient levels to allow them to utilize different dietary compounds more efficiently later in their life. Geurden et al. [1] first found that juvenile rainbow trout *Oncorhynchus mykiss* subjected to PP-based diet during the first three weeks of life showed improved acceptance and utilization of the same dietary PP-based diet when given during later life stages. Furthermore, two other studies reported that early exposure of young European seabass *Dicentrarchus labrax* to diets deficient in long-chain polyunsaturated fatty acids induced higher expression of delta-6 desaturase mRNA levels in juveniles – the rate-limiting enzyme involved in biosynthesis of highly unsaturated fatty acids [2, 3]. Other studies also point out the importance of maternal NP and its impact on the well-being of progeny. Izquierdo et al. [4] found that it is possible to achieve improved growth of 4-month old gilthead seabream *Sparus aurata* juveniles fed low FM and low fish oil diets by previously exposing the broodstock of those fish to high vegetable-based oil feeds. Moreover, these authors later showed that the same fish at 16-month of age were still able to grow on low FM/fish oil diets better compared to control group suggesting positive long-term effect of NP on utilization of vegetable-based diet [5]. Finally, NP with soybean meal-based diets has been recently shown to successfully adapt yellow perch *Perca flavenscens* to utilize the same soybean meal diet during adult stages, leading to better growth compared to yellow perch that were not exposed to soybean meal diet during the early developmental stage [6]. Most of the NP studies on fish focus, however, on NP with PP induced during fish juvenile stages. Perera and Yufera [7] attempted to nutritionally program zebrafish *Danio rerio* in its larval stage with soybean meal-based feeds at first feeding but no improvement in the growth of pre-adult fish was reported, likely due to poor dry feed intake at the NP stage. It is therefore critical to understand if an optimal timing exists for NP to take effect and improve the capacity of the fish to utilize PP for growth in later life stages. Zebrafish requires live food feeding during the first days of live for proper growth, development, and survival, and therefore NP of larval fish with dry feed poses a high risk of limited feed intake leading to possibly confounding results. We believe that live food can be used as a vector to induce NP during these very early developmental stages.

Although NP has a great potential to improve fish growth and health performance during the grow-out phase, the mechanism behind the NP phenomenon remains elusive. Flavor experiences during early life stages have been shown to be a driver for life-long flavor acceptance in adults’ life. For example, citric acid, which has a sour/bitter taste, is known to be an instinctively aversive substance for rats. However, rats exposed/programmed to citric acid during nursing showed increased voluntary ingestion of citric acid compared to rats that were not exposed to citric acid at all [8]. Similarly, Balasubramanian et al. [9] indicated that early PP diet exposure in rainbow trout might mediate feed acceptance of the same diet at a later life stage by affecting pathways regulating the sensory perception of taste, odor, and vision. If improved dietary PP utilization induced by NP is a result of improved palatability towards dietary PP, can the negative effects of anti-nutritional factors present in PP still induce morphological changes in fish digestive tract as seen in other studies [10–13]? Dietary PP, such as soybean meal, have been associated with many cases of intestinal inflammation in several fish species limiting their dietary inclusion rates [7, 10, 14–18]. Some of the typical signs of dietary PP-induced inflammation in the intestinal mucosa include shortening of the mucosal folds which decrease the capacity of the digestive tract to digest, absorb, and utilize nutrients consequently affecting fish growth and health by diminishing their response to pathogens [19]. Nutritional programming has been shown to improve dietary PP utilization, including soybean meal; whether this process is driven solely by increased palatability and therefore improved feed intake, or by specific intestinal responses to dietary PP that lead to morphological adaptations remains unknown. We believe that NP with dietary PP alters the gut epithelial lining and consequently increases fish resistance to negative side-effects of PP, thereby improving fish ability to cope with those alternative raw materials.

Fish gastrointestinal tract is characterized by certain amount of plasticity dependent on various signaling pathways which allow the gut to adapt to rapid shifts in environmental conditions including a diet [20]. Some of these signals are based on hormones that can be divided into two groups: those that induce (orexigenic), and those that inhibit (anorexigenic) appetite and food consumption. Peripheral signals originate in the gastrointestinal tract, liver, adipose tissue, and others, and reach the brain through both endocrine and neuroendocrine actions. The hypothalamus is considered the center that controls appetite and energy balance and integrates the different chemical (nutrient levels), mechanical (feed volume in the gut), and endocrine signals [21]. If improved taste is in fact the key driver for improved growth performance in previously “programmed” fish, we would expect to observe differential expression of digestive hormones responsible for appetite control after exposure to dietary PP between adult fish that were exposed to PP in their early age and those that were not. Therefore, the objectives of our study were: 1) To determine the effect of NP on dietary PP utilization in zebrafish; 2) To determine the effect of different NP induction times on dietary PP utilization; 3) To determine if NP induces any morphological changes in the gut responsible for improved digestion of dietary PP; and 4) To assess if NP with dietary PP affects the expression of appetite-stimulating hormones possibly responsible for enhanced dietary PP utilization.

The experiment was conducted with zebrafish since it is considered an established model species for various research areas including genetics, nutrition, biomedicine, and toxicology; additionally, its popularity is now growing in the aquaculture research field [22]. It is an omnivorous teleost species that can utilize both plant and animal protein sources efficiently for growth serving as a great model for assessment of nutritional pathways in carnivorous and herbivorous aquaculture species [22].

## Materials and Methods

The feeding trial was conducted in the Center for Fisheries, Aquaculture, and Aquatic Sciences at Southern Illinois University-Carbondale (SIUC), IL. All experiments were approved by the Institutional Animal Care and Use Committee (IACUC) of SIUC.

### Experimental fish

All experiments were carried using recirculated aquaculture system (Pentair Aquatic Ecosystems, Cary, NC) consisting of biofilter, carbon filter, UV light, and pH/conductivity automatic adjustment feature. The experimental culture system used reversed osmosis as the main water source where pH and conductivity were maintained at optimal levels of 6.9±0.2 and 1584±27 µS, respectively. The average water temperature throughout the feeding trial was 27.1 ± 0.2 °C. The photoperiod consisted of 14 hours of darkness and 10 hours of light, with the overhead lights on from 8:00-18:00.

Zebrafish broodstock was obtained from a local pet store (Petco, Carbondale, IL). Males and females were kept in separate tanks and fed 2-3 times a day with commercial feed (Otohime, Japan) and *Artemia* nauplii for two weeks before breeding. Fish were combined for breeding in a 2:1 ratio of females to males. A wire net with a 1.5 mm mesh was placed in breeding tank with artificial plants to induce spawning. The fish were left to breed for 24 hours. The broodstock was then removed after spawning on the next day. The wire mesh was taken out, and eggs hatched approximately 48 hours at 27 °C. At 3 days post-hatch (dph) after mouth opening and when the larvae started actively swimming, they were randomly distributed into experimental tanks.

Zebrafish were fed with rotifers *Brachionus plicatilis* only for the first two days after the swim up stage (3-4 dph). The rotifers were obtained from cysts purchased from a commercial vendor (Brine Shrimp Direct, Ogden, Utah). Starting at 5 dph all fish received *Artemia* nauplii obtained from hatching of cysts from a commercial source (GSL Brine Shrimp, Ogden, Utah) together with rotifers (5-7 dph) and then *Artemia* nauplii only from 8-13 dph. All groups were fed *ad libitum* throughout the study.

### Experimental diets

All the experimental feeds were formulated and produced at SIUC. Two experimental diets were tested: a diet based on soybean meal and soy protein concentrate as main protein sources replacing 80% of marine animal protein (PP diet; the soy concentrate was included to adjust dietary crude protein while leaving room for other ingredients in the formulation, including a minimum level of starch to allow expansion and floatability of the experimental diets); and a diet based on fishmeal as a main protein source (FM diet). Both diets were formulated to be isonitrogenous (49% crude protein) and isolipidic (10% lipid) (Table 1).

**Table 1.**
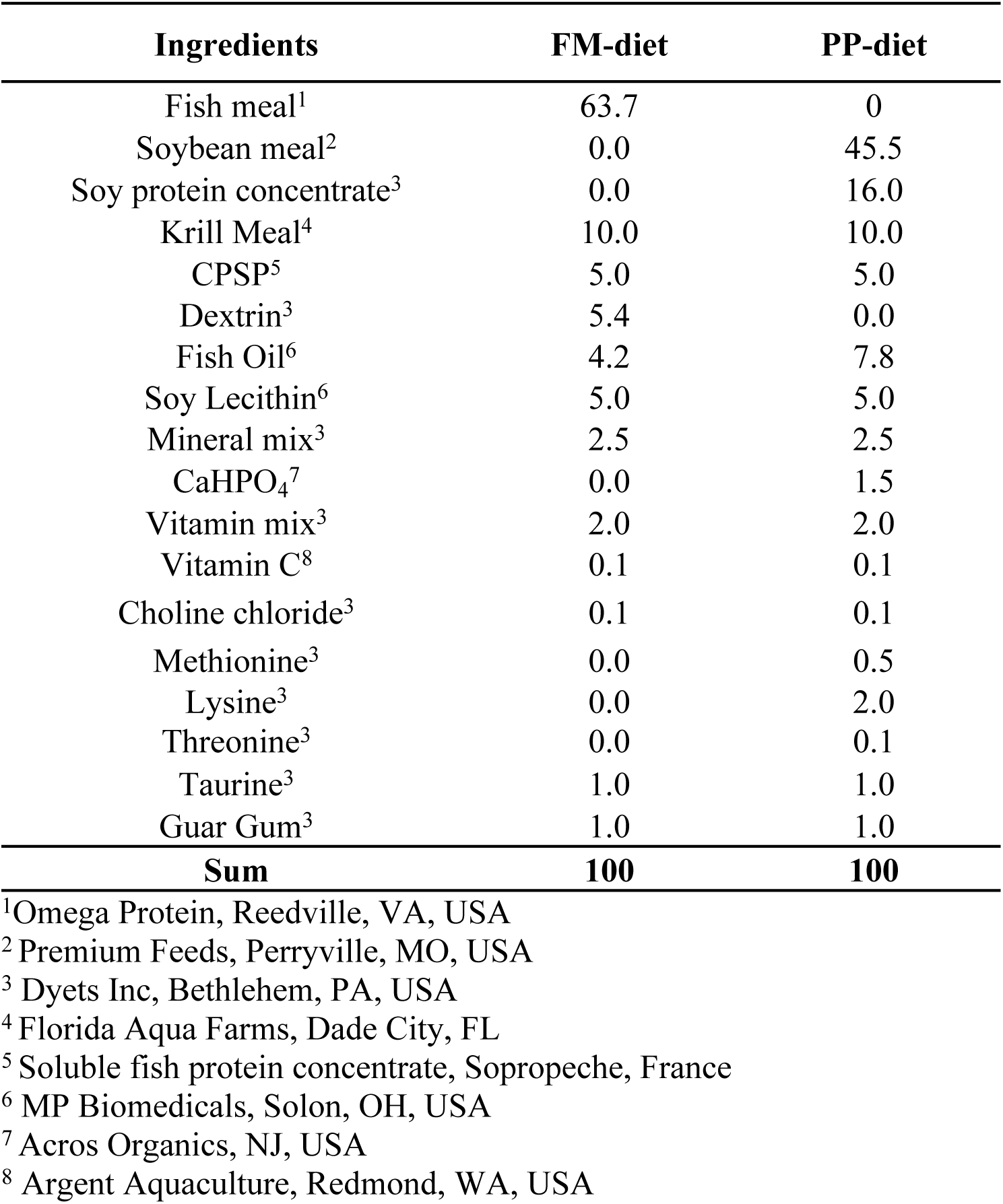
Dietary formulation (g/100 g) of experimental diet.

All dry protein ingredients (fishmeal, krill meal, and soybean meal) were added to a centrifugal mill (Retsch Haan, Germany) and ground to 0.5 micrometers. After the grinding process, all ingredients were manually sieved through a 0.25-micrometer sieve to ensure all particles were of the appropriate and uniform size. All the dry ingredients (excluding soy lecithin and choline chloride) were added together and mixed for 15 minutes and the fish oil was then added with the soy lecithin dissolved in the oil. The oil and dry ingredients were mixed again for 15 minutes. Finally water (∼10-15% of total mass of feed) was added with dissolved choline chloride. Feeds were then slowly added to the extruder (Caleva Extruder 20, Sturminster Newton Dorset, England) at levels between 20-24 rpm to obtain a proper extrudate size and firmness. Extrudates were then processed using a spheronizer (Caleva, Sturminster Newton Dorset, England) at 600 rpm for 3 min, 1800 rpm for 30 seconds, and then 600 rpm for 2-5 minutes to finish the process. Finally, the extrudates were dried using a freeze dryer (Labconco, Kansas City, MO). All dried pellets were sieved to appropriate sizes using a vibratory sieve shaker (Retsch Hann, Germany). All finished feeds were stored in bags at −20°C. While the feeds were being used in experimentation, they were kept at 4°C.

### Experimental design

At 3 dph zebrafish larvae were randomly distributed into 12 (3 L) tanks, 100 larvae per tank. The study lasted until fish reached 66 dph. Four different feeding regimes were investigated: 1) The first group received FM-based diet after the live food period from 13-36 dph and was challenged with PP-based diet between 36-66 dph (control, NP-FM); 2) The second group received NP with dietary PP in the early larval stage via live food enrichment during 3-13 dph followed by fishmeal (FM)-based diet during 13-36 dph and PP-based diet during 36-66 dph (NP-PP); 3) The third group received NP with dietary PP between 13-23 dph through formulated diet after the live food period followed by FM-based diet during 23-36 dph and PP-based diet during 36-66 dph (T-NP); and 4) The fourth group received PP-based diet after the live food period from 13-66 dph (negative control; PP) (Figure 1).

**Fig 1.**
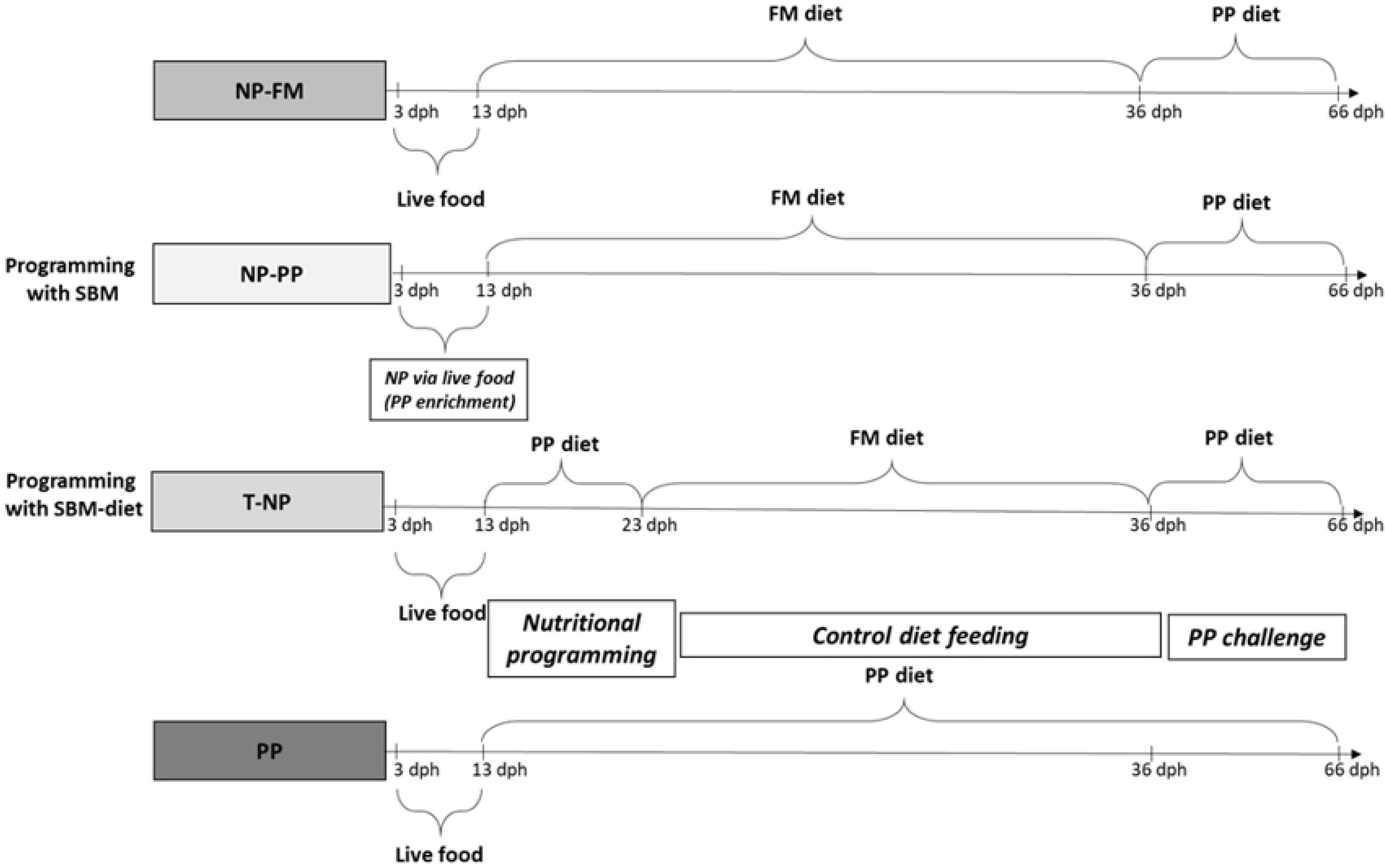
Dietary treatment regimens tested in the study. NP – nutritional programming, FM – fishmeal, PP – plant protein, SBM – soybean meal.

For the NP of the NP-PP group an enrichment of the live food was prepared with soybean meal. Both rotifers and *Artemia* nauplii were enriched by adding previously sieved finely ground soybean meal (<0.15mm) into the water of the rotifer or *Artemia* culture for a minimum of 3 hours prior to fish feeding (Figure 2). All the other groups (NP-FM, T-NP, and negative control) received live food enriched with Spirulina algae (Earthrise, Irvine, CA) using the same enrichment procedure.

**Fig 2.**
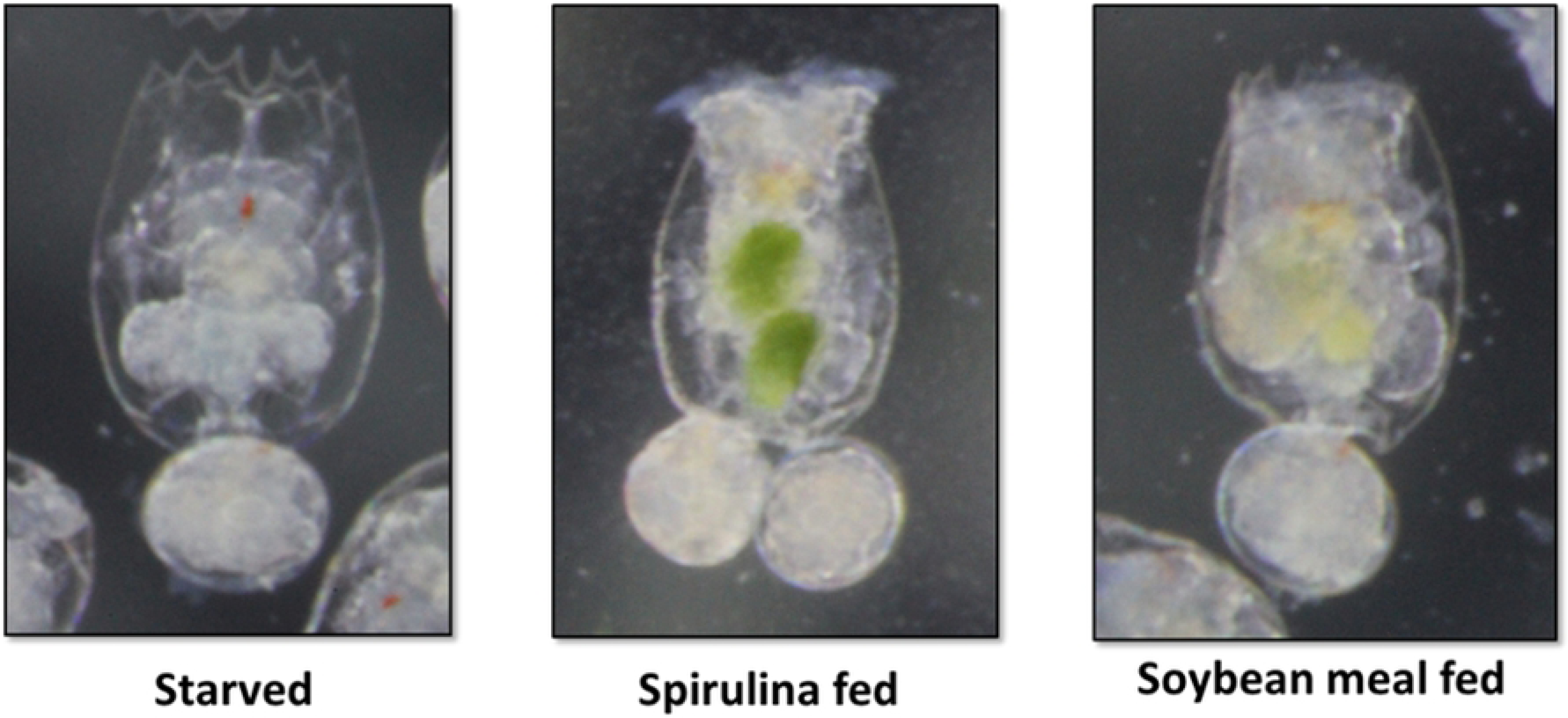
Rotifers after 2 hours of enrichment with Spirulina or ground, blended, and sieved soybean meal.

### Measured responses

At the end of the feeding trial, fish in each tank were counted and weighed. The following quantified growth performance parameters were assessed:

- Survival (%) = 100 × (final number of fish/initial number of fish)
- Final Weight (g) = Final body weight – initial body weight
- Weight gain (% of initial weight) = 100 × (final body weight – initial body weight)/initial body weight.

At the beginning (NP phase), in the middle (control phase), and at the end of the trial (PP challenge) feed intake was assessed by measuring the amount of feed that was consumed in one meal in each tank (feeding was ceased when fish showed signs of feed rejection).

At the end of the study, samples of fish were taken 3 hours (postprandial levels) and 24 hours (physiological baseline) after feeding from each group and preserved in RNALater® (Sigma-Aldrich, St Louis, MO) to assess the expression of digestive hormones. In addition, three fish from each tank were sampled, their digestive tracts were dissected and preserved in 10% buffered formalin for histological evaluation of the intestinal villi.

### Digestive hormone analysis

The digestive hormones gene expression analysis was conducted in the laboratory of Animal Biotechnology and Aquaculture of the Department of Biotechnologies and Life Sciences at University of Insubria, Varese, Italy.

### Total RNA extraction and cDNA synthesis

Total RNA was extracted from zebrafish brain and intestine samples using an automatic system (Maxwell® 16 Instrument, Promega) and a total RNA purification kit (Maxwell® 16 LEV simplyRNA Tissue). The extracted RNA was quantified by measuring the absorbance at 260 nm using NanoDrop™ 2000c spectrophotometer (Thermo Scientific) and reverse transcribed into cDNA following the protocol described in the SuperScript III Reverse Transcriptase kit (Invitrogen, Milan, Italy)

### Primer design and amplification of gene target

The primers for the amplification of *ghrelin, leptin, cholecystokinin (CCK), orexin*, and *neuropeptide Y (NPY)* genes were designed based on the cDNA sequences of these genes in *Danio rerio* available in GenBank database. The accession numbers of the sequences are **<u>AM055940.1</u>** for *ghrelin*, **<u>BN000830.1</u>** for *leptin*, **<u>XM_001346104.6</u>** for *CCK*, **<u>NM_001077392</u>** for *orexin*, and **<u>BC162071.1</u>** for *NPY* gene. The primer sequences are listed in Table 2.

**Table 2.**
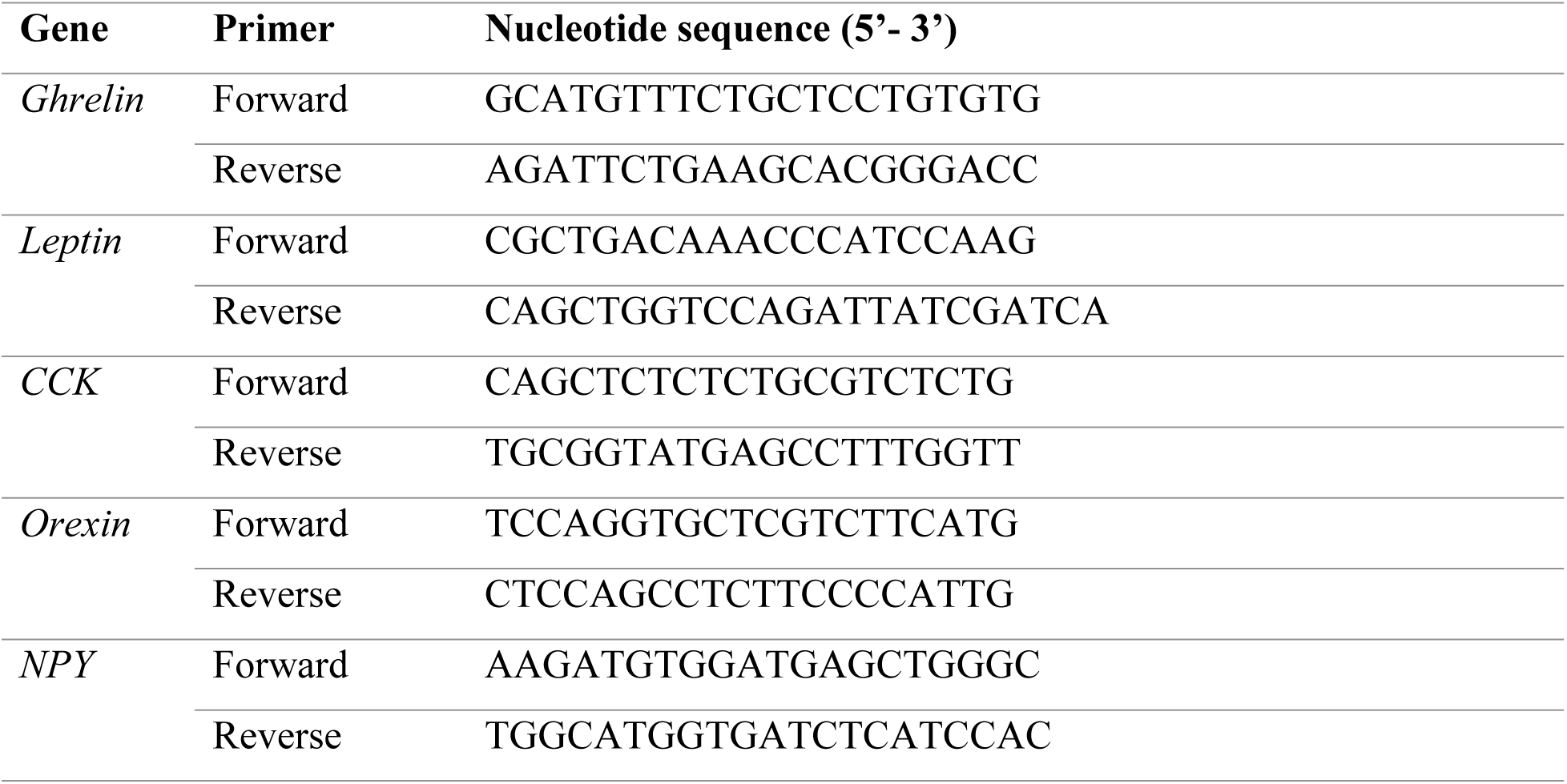
Primers used for the molecular cloning of target genes.

### Molecular cloning and sequencing

An aliquot of cDNA obtained by reverse transcription was amplified in a PCR reaction with GoTaq Green Master Mix (Promega, Milan, Italy). For each pair of primers, 25 PCR amplification cycles were set using the T100 Thermal Cycler (BioRad, Milan, Italy) thermocycler. The amplification product was then loaded onto 1% agarose gel with ethidium bromide in TAE 1X buffer to verify the length of amplicon. The bands of interest were extracted from the gel, and then purified using the NucleoSpin® Gel and PCR Clean-up kit (Macherey-Nagel Milan, Italy). For each target gene, a ligation reaction was performed between the pGEM®-T Easy plasmid (Promega Milan, Italy) and the amplicon purified. Briefly, to achieve optimal ligation efficiency, the necessary quantity of insert was calculated using the following equation:

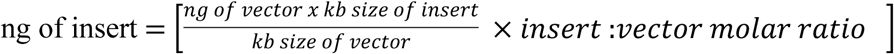

Then, following the protocol of the pGEM®-T and pGEM®-T kit, Easy Vector Systems (Promega, Italy), a mix was prepared containing: the insert, 5 μL of 2X Rapid Ligation Buffer, 1 μL of pGEM®-T Easy vector (50 ng), 1 μL of T4 DNA ligase (3U/μL) and water up to a final volume of 10 μL. The solution was left at 4°C overnight. The following day a transformation was carried out with highly competent E. coli JM109 (Promega) cells, which were added to the ligation solution, subjected to thermal shock (42 °C) for 45 seconds and allowed to grow in LB medium (Luria Broth) for 90 min.

Finally, the transformed cells were plated and grown overnight at 37°C. By means of white-blue screening it was possible to discriminate the colonies containing the vector, which were taken and subjected to a PCR reaction to check the presence of the insert of interest. Positive colonies were inoculated in liquid medium and left overnight in a shaker at 37 °C. The next day, the plasmid was purified using the NucleoSpin® Plasmid kit (Macherey-Nagel, Milan, Italy) from 4mL of culture, following the protocol provided by the manufacturer. The plasmids were evaluated on a 1% agarose gel and displayed on transilluminator (BioRad city, country). The amplification products were finally sequenced in both directions (T7 and SP6).

### Quantitative real-time RT-PCR

#### Generation of in vitro-transcribed mRNAs for target genes

Based on the cDNA sequences of zebrafish *ghrelin*, *leptin*, *CCK*, *orexin*, and *NPY* genes, sequenced as aforementioned, a forward and a reverse primer were designed for each target gene (Table 3). The forward primers were engineered to contain at the 5’ end the sequence of the T3 RNA polymerase promoter. The T3 RNA polymerase encoded by bacteriophages T3 is a DNA-dependent RNA polymerase that catalyzes the formation of RNA in the 5’→ 3’ direction after the viruses infect their host. The sequence of the T3 RNA polymerase promoter is necessary for the in vitro transcription of the mRNAs of each target gene to be used as standard in the subsequent quantification of the same gene in the biological samples. The T3 forward primers and their respective reverse primers were used in a conventional PCR reaction starting from plasmid. Then, the PCR product was checked on 1% agarose gel with ethidium bromide, purified with the NucleoSpin® Gel and PCR Clean-up kit (Macherey-Nagel, Italy) and subsequently sequenced.

**Table 3.**
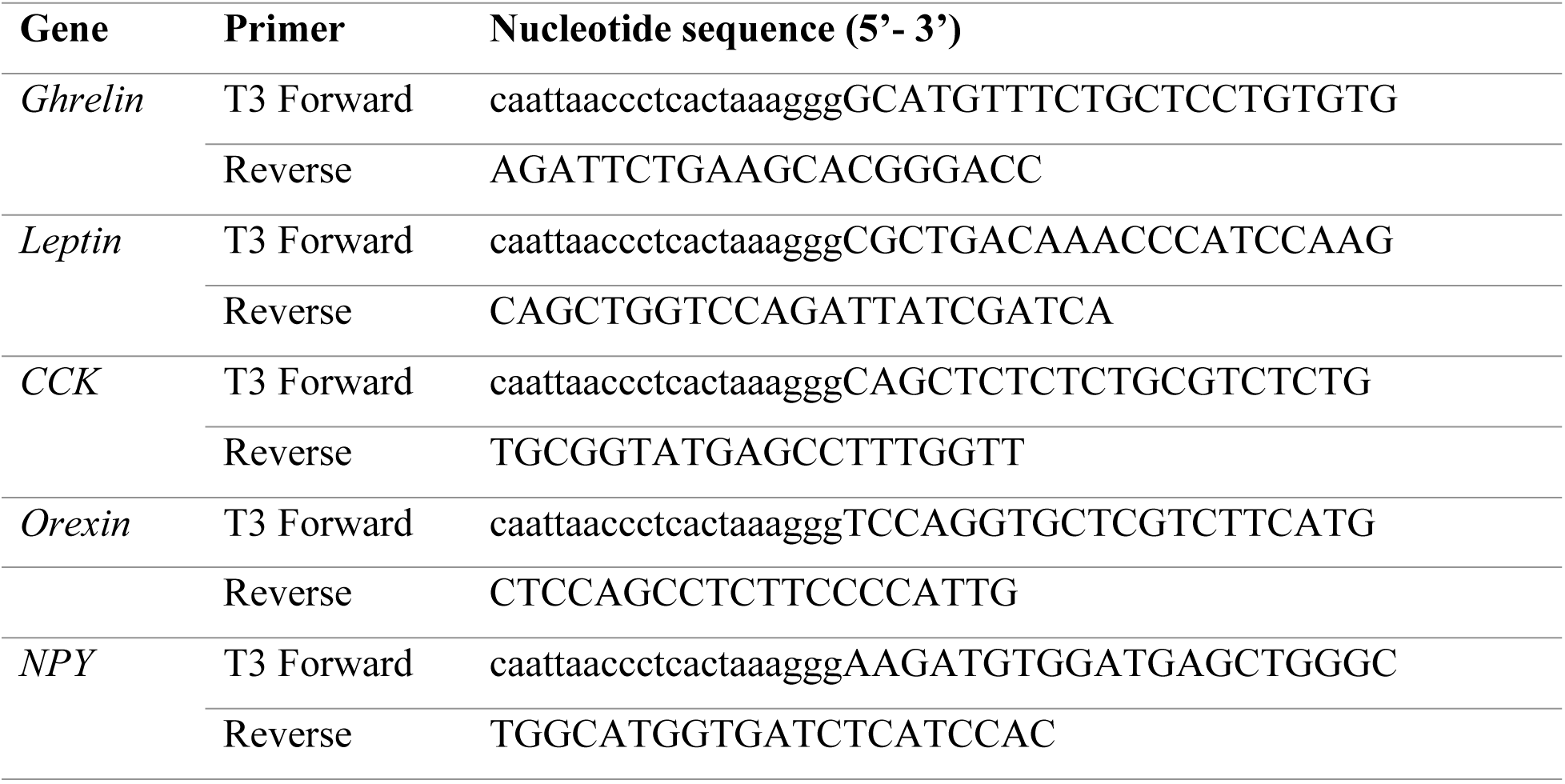
Primers used for the synthesis of standard mRNAs

Sequencing was necessary both to confirm the presence of the T7 promoter and to count the number of nucleotides present for the subsequent calculation of the molecular weight (MW - molecular weight) of each standard determined using the following formula: MW= [(n° of bases A x 329.2) + (n° of bases U x 306.2) + (n° of bases C x 305.2) + (n° of bases G x 345.2)] + 159.

In vitro transcription was performed using T3 RNA polymerase and other reagents supplied in the RiboProbe In Vitro Transcription System kit (Promega, Italy) according to the manufacturer’s protocol. The concentration of the mRNAs thus obtained was measured by reading the absorbance at 260 nm by means of NanoDrop ™ 2000c (Thermo Scientific). By knowing the molecular weight and the concentration of each mRNA it was possible to determine the number of molecules/μL for each target gene.

#### Generation of standard curves

The synthetic mRNAs of each gene were used as quantitative standards in the analysis of experimental samples. Defined amounts of mRNAs at 10-fold dilutions were amplified via real-time PCR using iTaq™ Universal SYBR® Green One-Step kit (Bio-Rad, Italy) using the following RT-PCR conditions: 10 min at 50°C (reverse transcription reaction), 1 min at 95°C, and 40 cycles consisting of 10 s at 95°C and 30 s at 60°C. The Ct values obtained by such amplification were used to create a standard curve for each of the target genes.

#### Real-time RT-PCR for quantification

One hundred nanograms of total RNA extracted from the experimental samples were amplified via One-step SYBR® Green quantitative real-time RT-PCR, in the same plate with 10-fold-diluited defined amounts of standard mRNA. The sequences of primers used for target gene amplification are shown in Table 4. SYBR® Green PCR reactions were performed on a Bio-Rad® CFX96™ System.

**Table 4.**
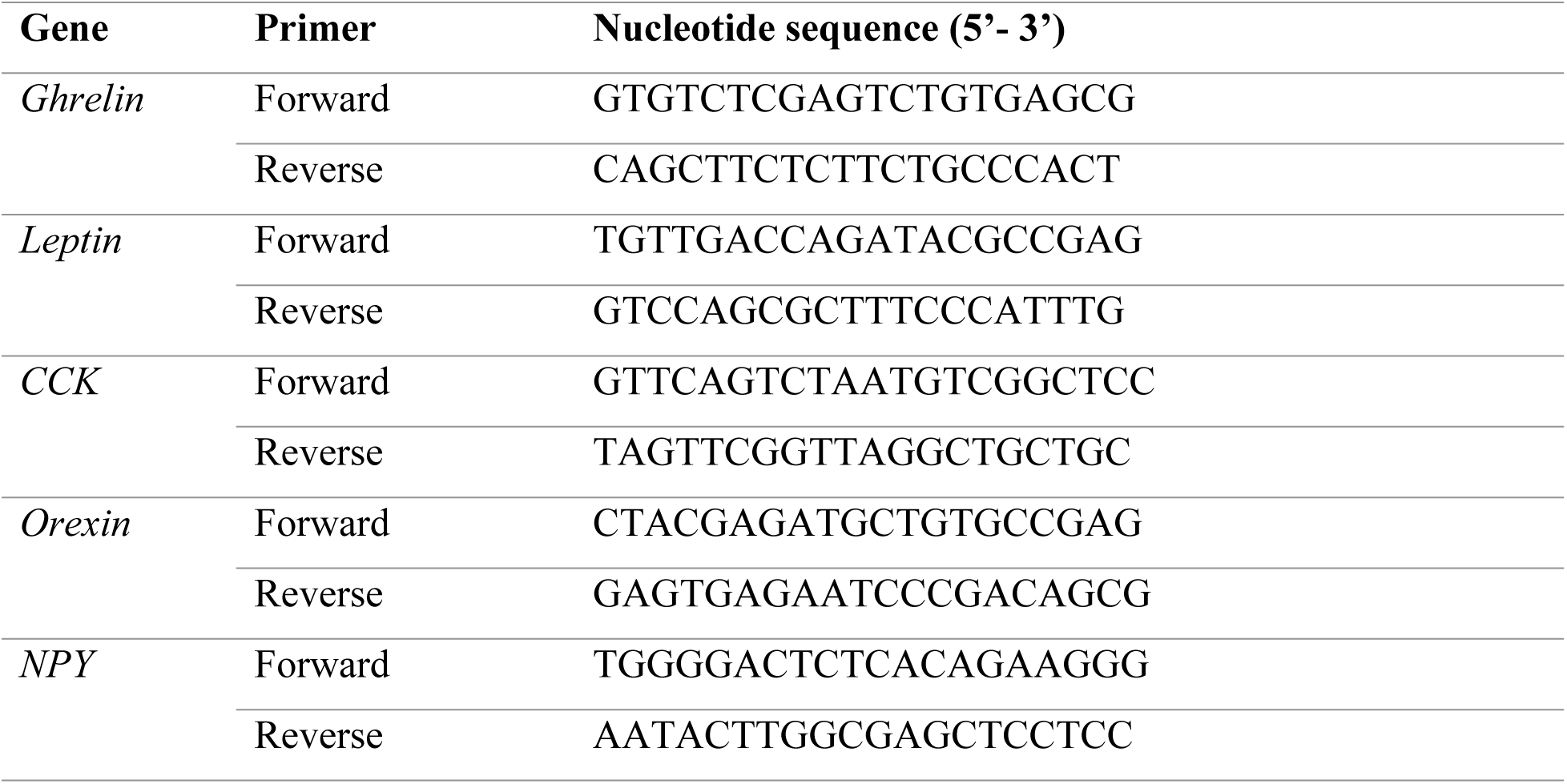
Primers and probes used for one-step SYBR^®^ Green real-time RT-PCR

Data from the real time PCR runs were collected with CFX™ Software. Cycle threshold (Ct) values corresponded to the number of cycles at which the fluorescence emission monitored in real time exceeded the threshold limit. The Ct values of the standard mRNAs amplification were used to create standard curves, which were then used for the calculation of the copies of mRNAs in the biologic samples.

### Histological analyses

The intestines previously fixed in 10% neutral buffered formalin were processed to paraffin using a Sakura enclosed automated tissue processor (Netherlands). The three representative areas of zebrafish intestines were orientated for cross sections embedded together in the same block. Five micrometer serial sections were cut with a Leica manual microtome (Buffalo Grove, IL) and placed on water bath at 44°C. Sections were placed on positive charged slides. After drying, the slides were stained with hematoxylin and eosin and cover-slipped using acrylic mounting media. The histological analysis of the mid-gut portions of fish digestive tract focused on the tissue sections of medium diameter. Pictures of the samples were taken at 100x magnification using a microscope (Nikon SMZ1500, Japan) and a camera (Nikon Digital Sight, Japan).

To obtain villus length and width, pictures of each slide were obtained using a microscope (Leica DMI 300B) and camera (Leica DMC 290) combination, with the software LAS V4.4 (Leica Camera, Wetzler, Germany). From these pictures, individual lengths and widths were taken of intact villi using ImageJ (NIH, Betheseda, MD, USA). Length and width data were measured according to Karimi and Zhandi [23] in the three different segments of the intestine; proximal, middle, and distal. Villi length was measured from the tip of the villus to the luminal surface, and villi width was measured across the base of the villus at the luminal surface. The length-to-width ratio of each villus was determined by dividing the length by the width.

### Statistical analyses

Results are presented as means ± standard deviation. One-way ANOVA was used followed by Tukey’s test using the software R. Differences with *p* values <.05 were considered significant.

## Results

### Growth performance

At the end of the study the lowest average weight was achieved by the PP group compared to all other groups. No differences were detected in average weight between NP-FM, NP-PP, or T-NP groups (Table 5).

**Table 5.** Dietary treatment effect on growth performance. Values are presented as means (± std. dev). Superscript letters indicate statistical significance between groups. The significance was determined using a One-Way ANOVA and a Tukey Test with a p value <0.05.

|  | NP-FM | NP-PP | T-NP | PP |
| --- | --- | --- | --- | --- |
| Avg. Weight (g) | 0.37<sup>a</sup> ($\pm 0.04$) | 0.36<sup>a</sup> ($\pm 0.05$) | 0.42<sup>a</sup> ($\pm 0.03$) | 0.26<sup>b</sup> ($\pm 0.03$) |
| Avg. Weight Gain (g) | 0.29<sup>a</sup> ($\pm 0.03$) | 0.28<sup>a</sup> ($\pm 0.03$) | 0.34<sup>a</sup> ($\pm 0.03$) | 0.20<sup>b</sup> ($\pm 0.03$) |
| Weight Gain (%) | 353.8<sup>ab</sup> ($\pm 14.7$) | 404.0<sup>ac</sup> ($\pm 45.1$) | 430.3<sup>c</sup> ($\pm 34.6$) | 364.6<sup>b</sup> ($\pm 29.9$) |

During the PP challenge the T-NP group achieved the highest weight gain compared to both control groups NP-FM and PP. The NP-PP group achieved higher weight gain compared to the negative control; however, no differences were detected with NP-FM or T-NP groups. No differences in the feed intake were detected during NP phase, control feeding phase, or PP challenge. Similarly, no differences were found in the survival (∼50% assessed for the trial duration from 3 until 66 dph) between the different dietary regimes.

### Digestive hormones expression

At the end of the study expression of digestive hormones was measured in the gut and/or brain 3 and 24-hours after feeding representing postprandial and basal levels, respectively. *Ghrelin* expression in the brain was significantly higher in the T-NP group compared to NP-FM and NP-PP groups but not different with the PP group. *Ghrelin* expression in the gut was significantly reduced in both NP-PP and T-NP groups compared to the other groups. No differences were detected in the gut *ghrelin* expression between NP-PP and T-NP nor between NP-FM and PP groups (Figure 3).

**Fig 3.**
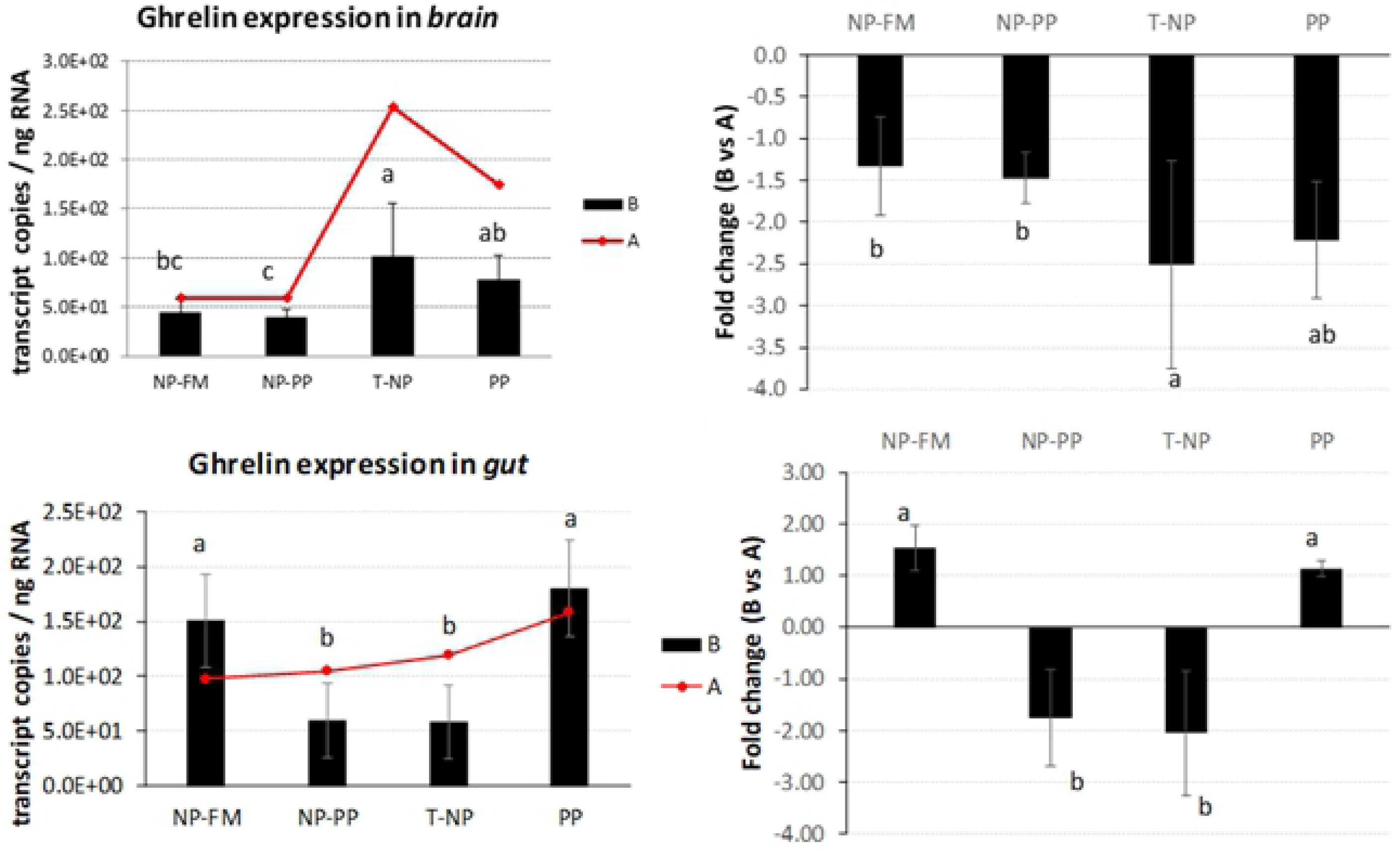
*Ghrelin* expression in the brain and gut. The expression of *ghrelin* presented as number of *ghrelin* transcript copies per ng of total RNA in zebrafish brain and gut at 3-hours (B) and 24-hours (A) after feeding. The fold change presents the difference in *ghrelin* transcript copies between 3 and 24-hour samples. Different letters indicate statistical difference at *p*<0.05.

*CCK* expression in the brain was significantly reduced in NP-PP and T-NP compared PP group but not different with NP-FM. *CCK* expression in the gut showed a similar trend, levels were lower in NP-PP and T-NP groups compared to PP but no difference was detected between T-NP and NP-FM (Figure 4).

**Fig 4.**
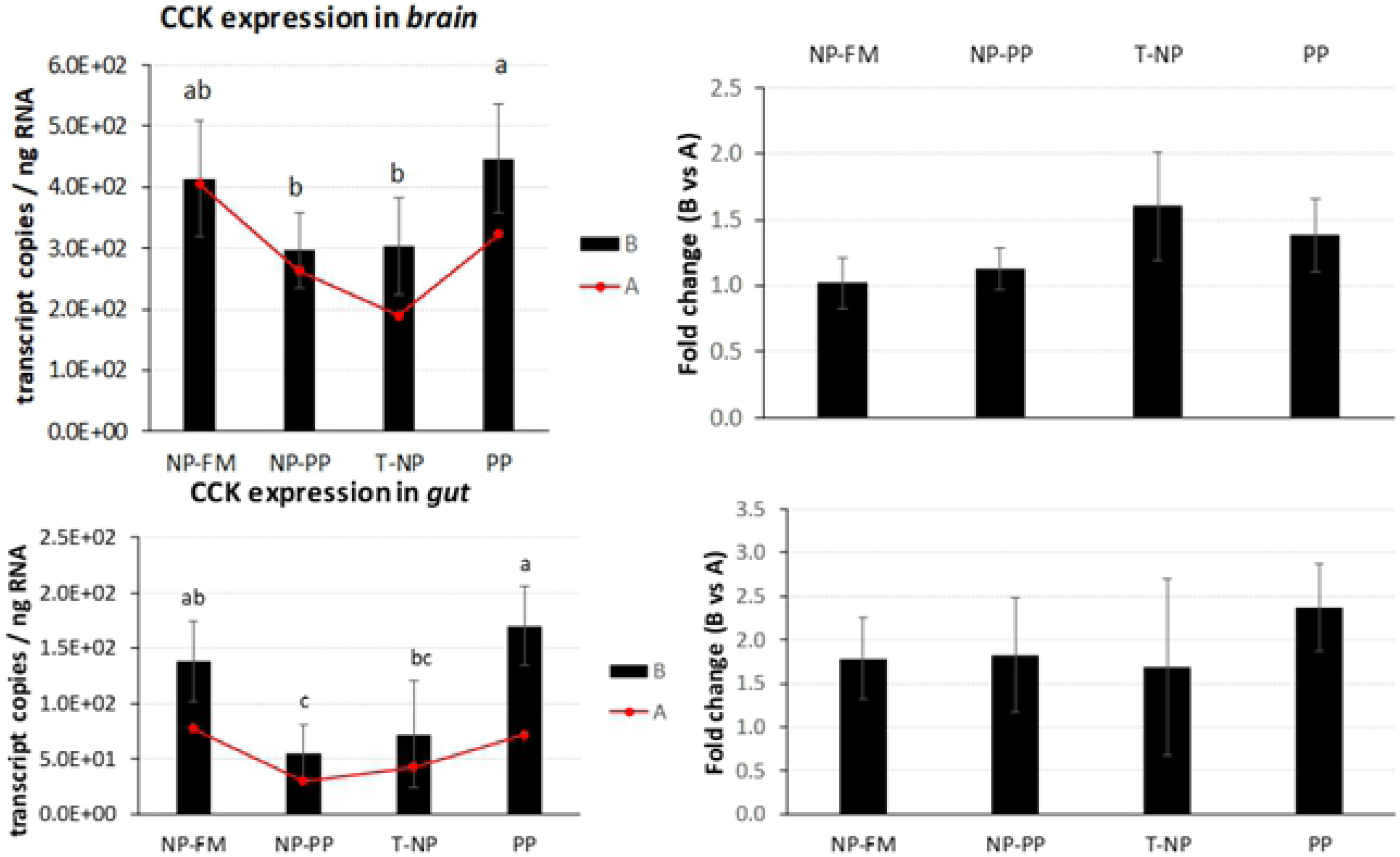
*CCK* expression in the brain and gut. The expression of *CCK* presented as number of *CCK* transcript copies per ng of total RNA in zebrafish brain and gut at 3-hours (B) and 24-hours (A) after feeding. The fold change presents the difference in *CCK* transcript copies between 3 and 24-hour samples. Different letters indicate statistical difference at *p* <0.05.

*Leptin* expression in the brain and gut was not different between groups 3-hours after feeding, but the level of *leptin* mRNA in T-NP fish 3-hours after feeding in the gut was lower compared to 24-hour sampled fish (Figure 5).

**Fig 5.**
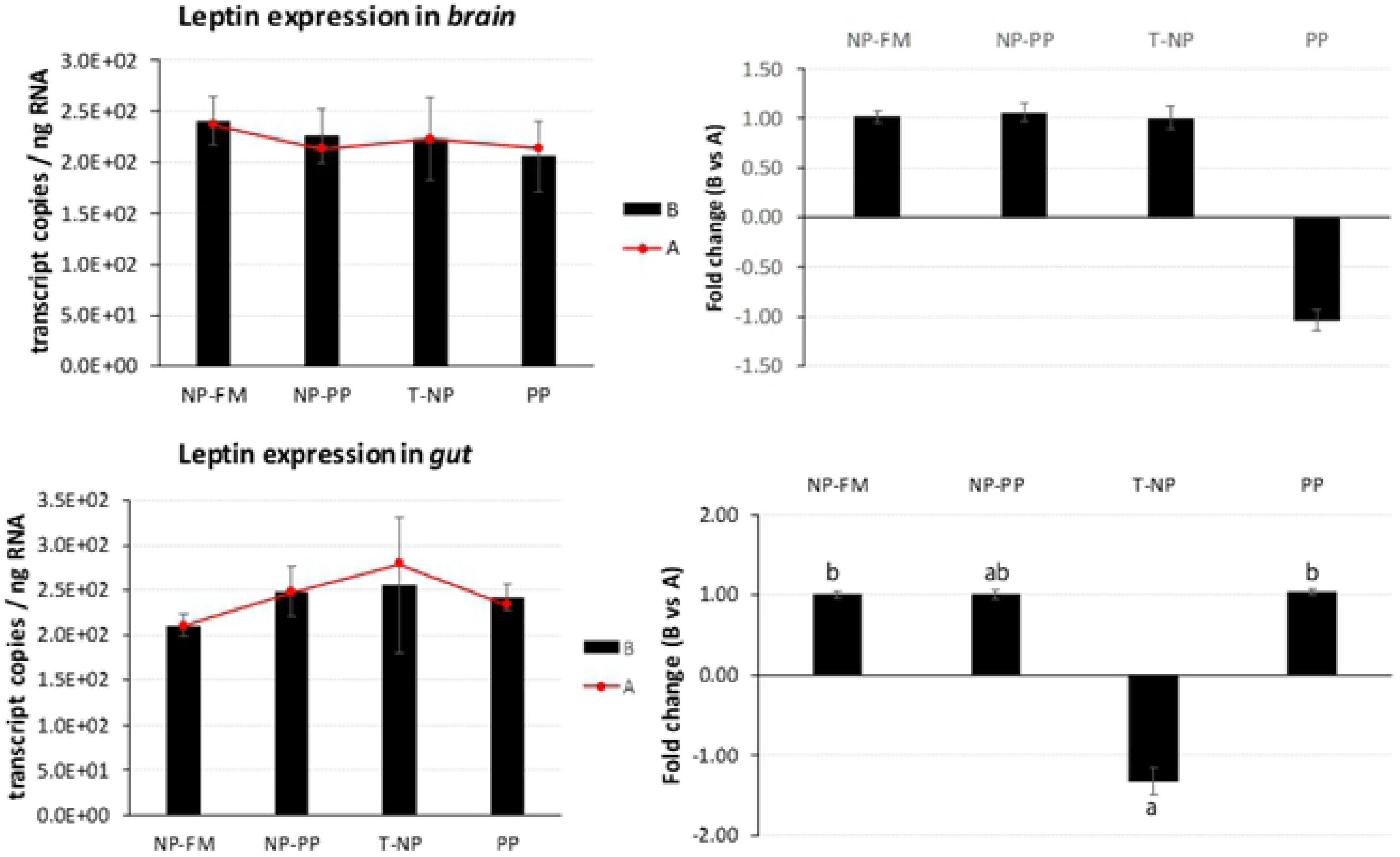
*Leptin* expression in the brain and gut. The expression of *leptin* presented as number of *leptin* transcript copies per ng of total RNA in zebrafish brain and gut at 3-hours (B) and 24-hours (A) after feeding. The fold change presents the difference in *leptin* transcript copies between 3 and 24-hour samples. Different letters indicate statistical difference at *p* <0.05.

No significance was detected in *orexin* expression in the brain nor the gut. However, an interesting trend was observed showing numerically higher expression level of brain *orexin* in T-NP and PP compared to the other groups (*p*>0.05). A similar trend was also seen in the gut but no significant differences were actually detected (Figure 6).

**Fig 6.**
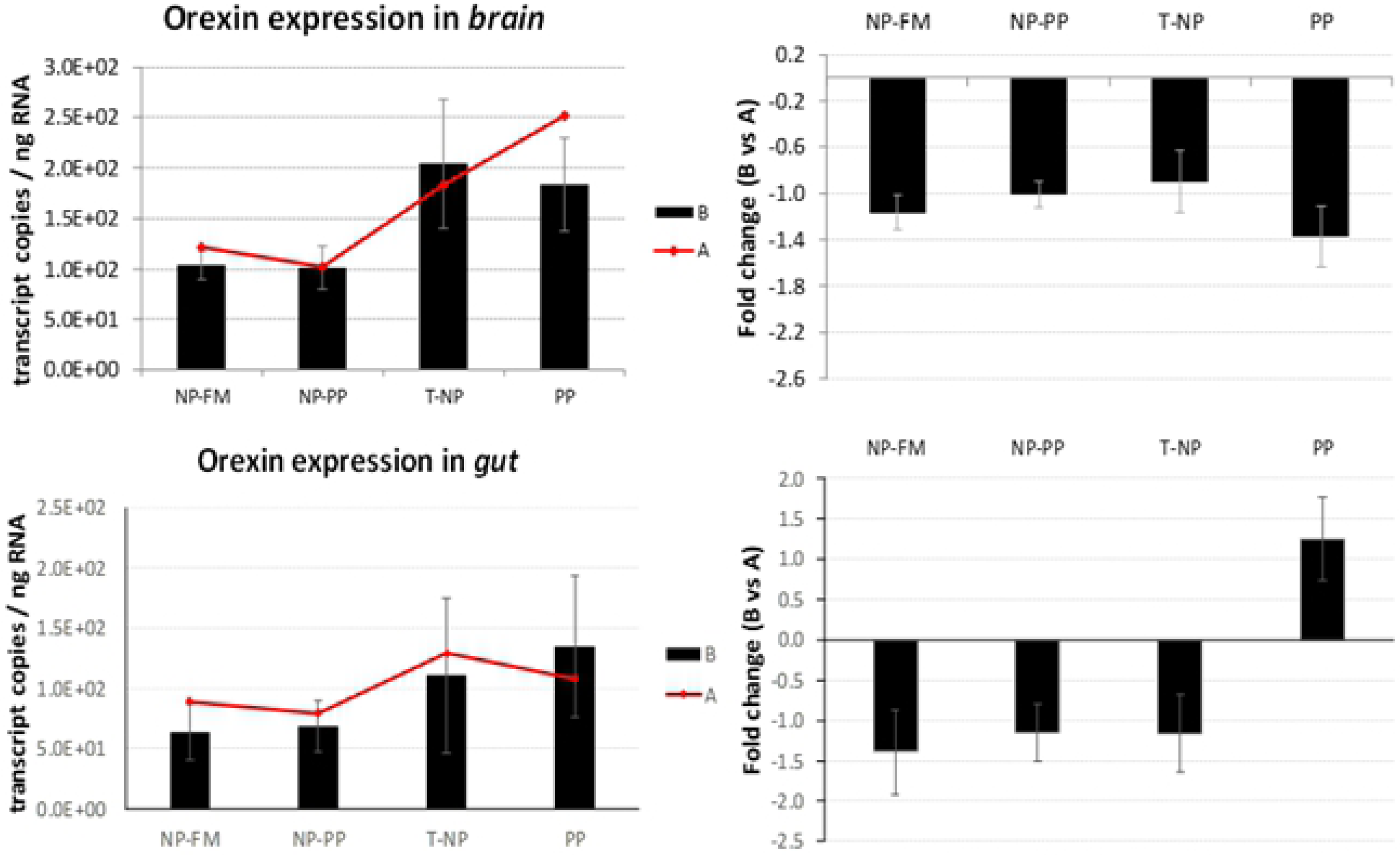
*Orexin* expression in the brain and gut. The expression of *orexin* presented as number of *orexin* transcript copies per ng of total RNA in zebrafish brain and gut at 3-hours (B) and 24-hours (A) after feeding. The fold change presents the difference in *orexin* transcript copies between 3 and 24-hour samples. Different letters indicate statistical difference at *p* <0.05.

*NPY* expression in the brain was significantly lower in the NP-PP compared to PP group but not different with NP-FM and T-NP groups (Figure 7).

**Fig 7.**
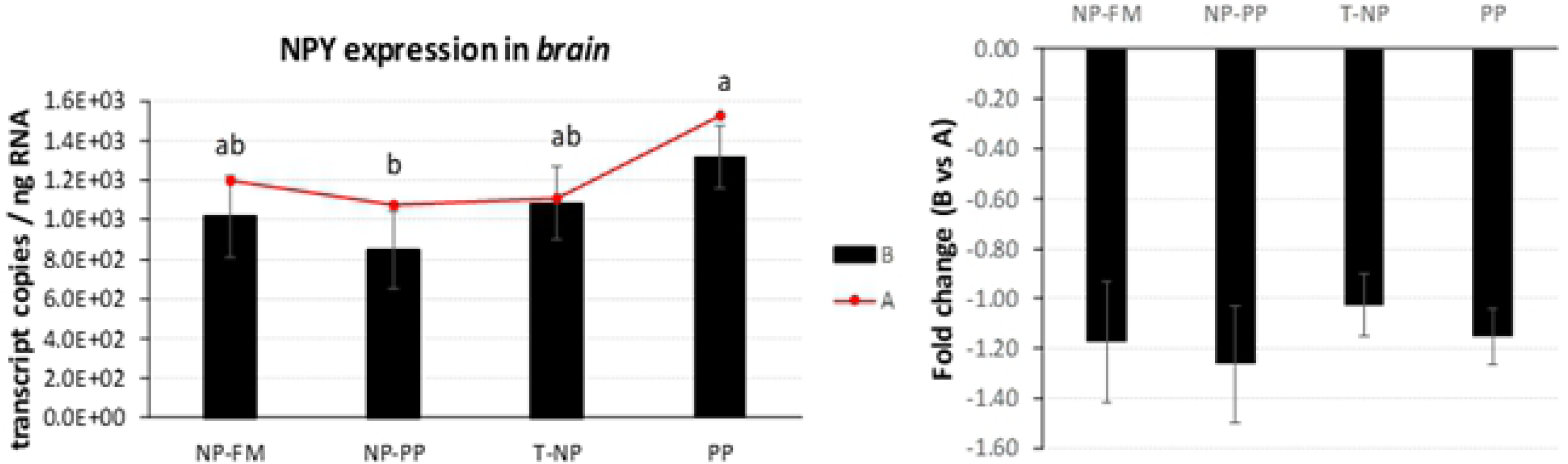
*NPY* expression in the brain. The expression of *NPY* presented as number of *NPY* transcript copies per ng RNA in zebrafish brain 3-hours (B) and 24-hours (A) after feeding. The fold change presents the difference in *NPY* transcript copies between 3 and 24-hour samples. Different letters indicate statistical difference at *p* <0.05.

### Intestinal villi

The length of intestinal villi in the proximal intestine was the highest in NP-FM group compared to all other groups. The villi width was found higher in NP-FM group compared to PP and T-NP groups but not different with NP-PP. In the middle intestine, however, the T-NP fish displayed higher villi length compared to PP group and not different compared to NP-FM and NP-PP groups. Finally, in the distal portion the T-NP group had the highest villi length compared to NP-PP and PP groups and not different with NP-FM group. No differences were detected in villi width in the middle or distal intestine.

The highest villus length to villus width ratio in the proximal intestine was found in T-NP compared to both NP-PP and PP groups and not different with NP-FM group. In the middle intestine the T-NP group had the highest villus length to width ratio compared to all other groups and no differences were detected between NP-FM, NP-PP, and PP groups. No differences were detected among groups in the length to villus ratio in the distal intestine (Table 6).

**Table 6.** Treatment effect on intestinal villi length, width, and length to width ratio. Units for villi length and width are µm. Values are presented as means (± std. dev). Superscript letters indicate statistical significance between groups. The significance was determined using a One-Way ANOVA and a Tukey Test with a *p* value <0.05.

| Intestinal Segment | Group |  |  |  |
| --- | --- | --- | --- | --- |
|  | NP-FM | NP-PP | T-NP | PP |
| **Proximal** |  |  |  |  |
| Length | 314.47<sup>b</sup> (± 58.14) | 213.12<sup>a</sup> (± 47.64) | 221.93<sup>a</sup> (± 72.21) | 188.24<sup>a</sup> (± 60.22) |
| Width | 112.18<sup>c</sup> (± 42.21) | 91.60<sup>bc</sup> (± 22.79) | 67.70<sup>a</sup> (± 13.40) | 79.43<sup>ab</sup> (± 17.80) |
| Ratio | 3.16<sup>ab</sup> (± 1.30) | 2.43<sup>a</sup> (± 0.66) | 3.37<sup>b</sup> (± 1.24) | 2.45<sup>a</sup> (± 0.91) |
| **Middle** |  |  |  |  |
| Length | 130.19<sup>ab</sup> (± 13.87) | 129.84<sup>ab</sup> (± 31.74) | 156.64<sup>b</sup> (± 33.71) | 121.97<sup>a</sup> (± 22.25) |
| Width | 70.09 (± 23.91) | 68.89 (± 22.86) | 56.92 (± 11.73) | 69.89 (± 14.98) |
| Ratio | 2.07<sup>a</sup> (± 0.78) | 1.99<sup>a</sup> (± 0.59) | 2.84<sup>b</sup> (± 0.72) | 1.80<sup>a</sup> (± 0.40) |
| **Distal** |  |  |  |  |
| Length | 160.92<sup>bc</sup> (± 21.99) | 106.41<sup>a</sup> (± 25.75) | 182.21<sup>c</sup> (± 43.33) | 130.62<sup>ab</sup> (± 52.30) |
| Width | 86.33 (± 20.76) | 63.63 (± 22.59) | 79.44 (± 20.18) | 81.86 (± 21.80) |
| Ratio | 1.98 (± 0.66) | 1.86 (± 0.82) | 2.40 (± 0.81) | 1.64 (± 0.71) |

## Discussion

Nutritional programming has been shown to improve growth performance in rainbow trout [1], yellow perch *Perca flavescens* [6], and gilthead seabream *Sparus aurata* [4, 5]. Perera and Yufera [7] induced NP during the first three days after mouth opening in zebrafish but did not observe any differences in the growth rate during later feeding with soybean meal-based diet. An induction of NP with dry feed as suggested by Perera and Yufera [7] in species that requires live food for proper growth, development, and survival, during the first feeding poses a risk of low or no food consumption defeating the actual NP outcome. We believe that programming with plant protein using a standard zebrafish rearing protocol utilizing live food as a vector for soybean meal could be a logical and straightforward approach in helping fish to adapt to this alternative protein source. Our experiment showed that dietary PP utilization and associated higher weight gain could potentially be improved in zebrafish with NP induced using live food as a PP vehicle and/or during zebrafish weaning period with dry PP-based feed. The experiment showed that after the PP challenge the T-NP group achieved the highest weight gain compared to both control groups suggesting that NP with dietary PP improves its utilization in zebrafish. The NP-PP group achieved higher weight gain compared to the negative control, however, no differences were detected with NP-FM or T-NP groups. It is possible that if the study was prolonged differences could also be detected between NP-PP and NP-FM groups. However, since the fish reach sexual maturation after approximately two months after hatching leading to size discrepancies within tanks and groups, the feeding trial had to be terminated at 66 dph before an obvious sexual dimorphism started to appear. Another possible scenario is that perhaps an optimal window exists for fish to respond to NP effectively. The live food used in the study, rotifers and *Artemia* nauplii, were both enriched with soybean meal and their enrichment status was assessed by change in body color (Figure 2) and therefore we believe there was an exposure to PP in larval zebrafish during the first days of feeding. It is possible, however, that fish must reach a certain developmental stage to be able to positively respond to the imposed nutritional trigger (such as soybean meal) to better adapt to it at a later age.

The available data on hormonal regulation of the gastrointestinal tract in teleosts suggest that some gastrointestinal hormones not only regulate digestion but also act as appetite/satiety modulating signals in the brain. When *ghrelin* was first discovered in rats it was demonstrated to be a powerful stimulator of food intake (the first known peripheral hormone with this effect) that stimulates body weight gain and adiposity in mammals. In the present study, fish brain and whole digestive tract samples were taken 3- and 24-hours after feeding. In T-NP fish 3-hours after feeding the level of *ghrelin* in the brain was higher compared to NP-PP and control fish. In the gut however, the level of *ghrelin* in both “programmed” groups was lower compared to control (NP-FM) and PP groups. Moreover, the level of *ghrelin* mRNA expression in NP-PP and T-NP groups 3-hours after feeding was lower compared to fish sampled 24-hours after feeding. *Ghrelin* in zebrafish is mainly expressed in endocrine pancreatic cells [24] and its higher mRNA levels have been associated with fasting [25]. The lower *ghrelin* expression level in the gut observed in our study might possibly indicate fish in “programmed” (NP-PP and T-NP) groups to be in a satiated (fed) state compared to both controls. In rainbow trout *Oncorhynchus mykiss ghrelin* was shown to have an anorexigenic effect that leads to suppression of food intake. However, in cyprinids such as goldfish *Carassius auratus*, *ghrelin* was reported to increase food intake through stimulation of *NPY* and *orexin* neurons in the brain and a potential feedback of *orexin* on *ghrelin* expression, which might help explain the elevated *ghrelin* levels in the T-NP fish in zebrafish brain in our study. *Orexin* has been found to stimulate appetite and food consumption after fasting in different fish species including zebrafish [26–29]. We found *orexin* expression levels in both brain and gut not significantly different among groups. However, there was a trend for higher levels of *orexin* in T-NP and PP groups compared to the other groups, particularly in the brain, possibly supporting the aforementioned assumption about the *orexin*-related increased *ghrelin* levels in T-NP group in the brain.

The role of *NPY* in feed regulation in fish has not been studied extensively but the available literature indicates that *NPY* in fish has orexigenic effects similar to those observed in mammals. More specifically, higher levels of *NPY* have been associated with increased food intake in zebrafish [30]. The *NPY* brain expression levels have been reported higher around feeding time and lower post-prandially in Chinook salmon *Oncorhynchus tshawytscha*, Coho salmon *Oncorhynchus kisutchand* [31], and in goldfish [32, 33] further suggesting a role of *NPY* as an appetite stimulator consequently being widely considered as one of the most highly conserved neuropeptides in vertebrates [30]. Our study found *NPY* expression to be higher in the PP group compared to NP-PP and not different with T-NP and NP-FM groups, suggesting that perhaps fish in the PP group were in an “unfed” state 3-hours after feeding. Valen et al. [34] observed higher *NPY* mRNA levels in fed Atlantic salmon *Salmo salar* compared to unfed fish 1.5 and 9-hours after feeding. The authors argued, however, that the increased *NPY* expression was probably a result of digestive tract feedback since after 9 hours the feed had likely been evacuated from the digestive tract rendering the fish to be in “unfed” state again.

*Leptin* - an antagonist of *ghrelin*, is an anorexigenic hormone released mostly by adipose tissue in mammals. In fish, *leptin* has been found in liver, brain, muscle, gonads, kidneys, and other tissues [21, 35–37] and its effects on energy metabolism in fish are still perplexing. An increased postprandial hepatic *leptin* expression have been observed in common carp *Cyprinus carpio* [36] and orange spotted grouper *Epinephelus coioides* [38] confirming its role as a satiety signal comparable to mammals. Murashita et al. [39] showed reduced growth performance of juvenile Atlantic salmon after intraperitoneal infusion of recombinant salmon specific *leptin* at a dose of 10 ng/g/h using implanted micro-osmotic pump over a three-week period. However, *leptin* response to feeding status varies between species and salmonids show an opposite trend to the mammalian model in which fasting or feed restriction actually leads to decreased *leptin* levels. Trombley et al. [37] reported 9-fold higher expression levels of hepatic *leptin* and 2.3 times higher plasma *leptin* levels in Atlantic salmon in the feed-restricted group (40% of the optimal feeding rate) compared to the control group (optimal feeding rate). Similar results were obtained by Kling et al. [40] who found plasma *leptin* levels to be higher in fasted rainbow trout compared to fed trout throughout 21 days of the experimental period. In our study, *leptin* expression in the brain was not significantly different among groups. *Leptin* expression in the gut was not different between groups 3-hours after feeding suggesting that either a longer time is required for the *leptin* signal to take effect or the link between diet and *leptin* regulation in zebrafish is not significant if all the fish receive similar food rations across groups as opposed to feeding versus starving. This first assumption can be consolidated by the observed rise in *leptin* mRNA levels in T-NP group 24-hours after feeding compared to 3-hour sampled fish potentially suggesting fish in T-NP group to be in a satiated state 24-hours after the last meal compared to all other groups. The second assumption is supported by results obtained by Oka et al. [41] who found no differences in *leptin* mRNA expression between overfed and control (optimal feeding rate) zebrafish. Huising et al. [36] reported that hepatic *leptin* expression levels corresponded to increase in plasma glucose levels 3-hours and 6-hours after feeding in common carp *Cyprinus carpio*. Yet, *leptin* expression was not affected by short-term fasting (six days) nor long-term-fasting (21 days) even though the latter fish lost 30% of their initial weight towards the end of the third week. *Leptin*’s role in satiety is supplemented by the actions of both *ghrelin* and *CCK*. Volkoff et al. [42] found that *CCK* might affect actions of *leptin* in goldfish. Interestingly, although in our study no differences were detected in *leptin* expression level among groups 3-hours after feeding, *CCK* expression level was significantly lower in both programmed groups compared to control and negative control in the brain. Similar trend was observed in the gut. *CCK* has been linked to stimulation of pancreatic enzyme secretion, gallbladder contraction, and intestinal peristalsis. In the brain *CCK* acts as a neurotransmitter involved in control of appetite as a satiation signal from the gut. In goldfish, *CCK* has been found to inhibit feed intake [43]. Consequently, decreased *CCK* expression in programmed groups potentially led to enhanced digestion and nutrient absorption and therefore better growth of fish in those two groups. However, *CCK* expression in the gut seemed to present a similar trend to *ghrelin* expression contradicting previous evidence on involvement of *ghrelin* in suppression of *CCK* anorexic effect in goldfish [44]. Although the majority of studies on several fish species indicate that fasting decreases *CCK* expression in the gut [45–50] the role of *CCK* response to fasting/feeding is species-, tissue-, and time-dependent [51]. For example, Macdonald and Volkoff [52] showed an increase in *CCK* expression in the gut after two weeks of fasting in winter skate *Raja ocellata* but no effect was detected in the brain. *CCK* has also been shown to differ between gastric and agastric fish species such as ballan wrasse *Labrus bergylta* [53]. The decreased expression of *CCK* in NP-PP and T-NP groups could also indicate these fish were actually in “starvation” compared to both control groups - an alternative scenario to the earlier statement on *CCK* status in our experiment. However, this speculation contradicts previous results presented for *ghrelin*, *orexin*, *NPY*, and *leptin* and therefore the results remain perplexing and suggest further investigation on the effects of nutritional status on the expression of *CCK* in zebrafish.

Soybean meal is the most commonly used PP source in aquaculture feeds, but its use has been limited due to its negative effects at higher dietary inclusion rates that lead to reduced growth and an intestinal inflammation [54]. Typical signs of dietary PP-induced inflammation in the intestinal mucosa include thickening of lamina propria and sub-epithelial mucosa; infiltration of inflammatory cells; and increased number of goblet cells in the epithelium; as well as shortening of the mucosal folds and hence, reduced absorptive capacity of the enterocytes lining the intestinal epithelium [19]. An increase in intestinal villi length has been associated with higher absorption of available nutrients due to increased surface area. However, assessment of villi length only does not consider variations in the surface area due to changes in intestinal villi width and consequently a ratio of intestinal villus length to width is commonly used. Our histological results revealed that the highest villus length to villus width ratio in the proximal intestine was found in T-NP and NP-FM groups compared to both NP-PP and PP groups. However, in the middle intestine the T-NP group had the highest villus length to villus width ratio compared to all other groups. Based on the differences in morphology and cell differentiation and function, Ng et al. [55] proposed that zebrafish intestinal tract should be divided into three distinct segments: the intestinal bulb (proximal intestine), middle intestine, and distal intestine. The digestive processes mostly occur in proximal and middle segments where absorptive enterocytes, digestive enzymes, and solute transporters are highly concentrated, while the distal region is likely involved in water and iron absorption [55, 56]. Our results could therefore suggest that increased villus length to width ratio in T-NP group could be a result of morphological adaptation of the intestine allowing for more efficient dietary soybean meal digestion and nutrient absorption. Wang et al. [16] reported that 100% dietary fishmeal replacement with soybean meal led to inflammatory response manifested by significant reduction in intestinal villi length, width, and length to width ratio similar to the results obtained in our negative control group. The intestinal inflammation caused by presence of anti-nutritional factors in dietary soybean meal has been studied in both carnivorous and omnivorous species [14–16, 57, 58] where zebrafish has been suggested as a model species due to presence of similar inflammatory responses after ingestion of different plant-based ingredients [10].

## Conclusions

Our study found that zebrafish is able to utilize PP-based diets more efficiently for growth when exposed to the same PP source early in life. This study also proposes that NP should probably be induced when fish are already in a juvenile stage when all the organs and systems are fully present to be able to respond to this early nutritional trigger more effectively. Finally, our study suggests that the mechanism behind NP might be associated with endocrine and morphological adaptation of the digestive system that leads to enhanced digestion and absorption capacity ultimately reflected by improved growth.

## Acknowledgments

We thank Saffron Scientific Histology Services (Carbondale, IL) for providing excellent histological services in a timely manner.

## Author contributions

Experimental conception and design: KK MW. Experiment management and execution: MW KK. Data analyses: GT, VJM, FI, GSM. Writing (original draft): KK. Writing (review and edit): KK MW GT VJM GSM.

